# Posture-dependent modulation of marmoset cortical motor maps detected via rapid multichannel epidural stimulation

**DOI:** 10.1101/2024.01.23.576825

**Authors:** Mitsuaki Takemi, Banty Tia, Akito Kosugi, Elisa Castagnola, Alberto Ansaldo, Davide Ricci, Luciano Fadiga, Junichi Ushiba, Atsushi Iriki

## Abstract

In this study, rapid topographical changes were detected in the forelimb motor maps in the primary motor cortex (M1) of awake marmoset monkeys using our previously developed accurate short-time stimulation mapping procedure (Takemi et al. 2017; Kosugi et al. 2018). The results revealed that although the hotspot (the location in M1 that elicited a forelimb muscle twitch with the lowest stimulus intensity) remained constant across postures, the stimulus intensity required to elicit the forelimb muscle twitch in the perihotspot region and the size of motor representations were posture-dependent. Hindlimb posture was particularly effective in inducing these modulations. The angle of the body axis relative to the gravitational vertical line did not alter the motor maps. These results provide a proof of concept that a rapid stimulation mapping system with chronically implanted cortical electrodes can capture the dynamic regulation of forelimb motor maps in natural conditions. The flexible nature of the motor maps necessitates the reconsideration of the results of motor control and neuroplasticity studies. Neural mechanisms regulating forelimb muscle representations in M1 by the hindlimb sensorimotor state warrant further exploration.

## Introduction

The primary motor cortex (M1) contains anatomically contiguous body muscle representations arrayed along contiguous regions of the cortical surface, forming a somatotopic map. However, this somatotopic organization is not as segregated as indicated by the classical motor homunculus (Penfield and Boldrey 1937). Evidence has revealed that the motor representations of the digits, wrist, forearm, and proximal arm are partially overlapping (Schieber and Hibbard 1993; Catani 2017) and can be adaptively altered via extensive hand motor training (Nudo et al. 1996a, 1996b). Furthermore, the topographical pattern of motor representations can change within minutes following motor nerve injury (Sanes et al. 1988) owing to the recruitment of existing neural pathways that are normally inhibited (Jacobs and Donoghue 1991).

The topography of M1 motor maps may promptly change depending on peripheral sensory inputs, which can alter GABAergic inhibition in the M1 (Ridding and Rothwell 1999). Further, single-neuron recordings in monkeys have revealed that the preferred movement direction of individual M1 neurons innervating forelimb muscles can be altered instantaneously by a change in static arm posture (Scott and Kalaska 1995; Kakei et al. 1999). These observations indicate that sensorimotor-driven dynamic adaptations in forelimb motor representations arise because of changes at the single-neuron level. Extending this notion to topographical motor representations, we recently reported that tactile brushing stimuli applied to rodent forelimbs induced a rapid expansion of the M1 forelimb representation (Kosugi et al. 2019). Additionally, the results revealed that this expansion persists for several minutes following the completion of the skin stimulation, thus indicating the dynamic nature of M1 motor maps (Kosugi et al. 2019). However, it is unclear whether these dynamic M1 maps change during continuous shifts in natural sensorimotor inputs, such as changes in whole-body posture, mostly due to technical limitations.

Unlike the common approach of modifying the sensory inputs directly into the forelimb, the current study uniquely explored the impact of the sensorimotor state of other body segments on the topographical motor representation of the forelimb muscles. To assess the dynamic aspect of M1 motor maps, we used a previously established fast and accurate motor mapping system (Takemi et al. 2017; Kosugi et al. 2018). This system minimizes temporal effects on the mapping results and can detect slight alterations in motor maps. We selected common marmoset monkeys as the animal model and a minimally invasive cortical surface stimulation (CSS) for M1 mapping method. The common marmoset has a nearly lissencephalic cortex with considerably thinner dura mater compared with that of the macaque monkey (Bourne and Rosa 2003). This structural characteristic enabled us to perform M1 mapping via CSS using epidural microelectrocorticographic (µECoG) electrodes. CSS causes less damage to the neurons than intracortical microstimulation (ICMS), which requires inserting needle electrodes into the brain (Rajan et al. 2015). Therefore, CSS can be suitable for longitudinal studies that involve repeated M1 mapping. The small body size of marmosets renders them easy to handle and suitable for whole-body posture manipulation.

Herein, we constructed M1 forelimb maps under three different whole-body postures: horizontal (trunk fixed to a horizontal pole and four limbs touching the pole), vertical (trunk fixed to a 60° inclined pole and four limbs touching the pole), and vertical-sitting (similar to the vertical posture, except that the hindlimbs were touching the ground; Fig. 2A). These postures could dissociate two sensorimotor factors that affect the motor maps: 1) static visual inputs and vestibular information against gravity and 2) the sensorimotor state of the hindlimbs. Moreover, these postures reflect natural ethological conditions. Given that marmosets use diverse body postures and occupy large three-dimensional volumes (Ngo et al. 2022), this study addresses how postural information is integrated with forelimb motor control in this species.

## Materials and Methods

### Animals

This study was conducted on three adult male marmosets (*Callithrix jacchus*; MK1, 358 g; MK2, 356 g; MK3, 380 g). Marmosets MK1 and MK2 were also included in a study that examined grasping- and locomotion-related cortical activities (Tia et al. 2017, 2021). Cortical mapping data in the horizontal posture were reported in our previous study (Kosugi et al. 2018), which aimed to establish a reliable long-term mapping algorithm for marmosets. All experimental procedures were performed according to the guidelines of the Laboratory Animal Welfare Act and the Guide for the Care and Use of Laboratory Animals (National Institutes of Health, Bethesda, MD). The study protocols were approved by the Institutional Animal Research Committee at RIKEN (IRB approval number H24-2-228 & H26-2-211).

### Microelectrocorticographic electrode arrays

Mapping of M1 was conducted by electrical CSS using an 8 × 8 µECoG electrode array sheet developed in our laboratory (Fig. 1A; Tia et al. 2017). The electrodes were coated with a nanocomposite of poly-(3,4-ethylene-dioxythiophene) and carbon nanotubes and encapsulated by fibrin hydrogel (Castagnola et al. 2013).

**Figure 1.**
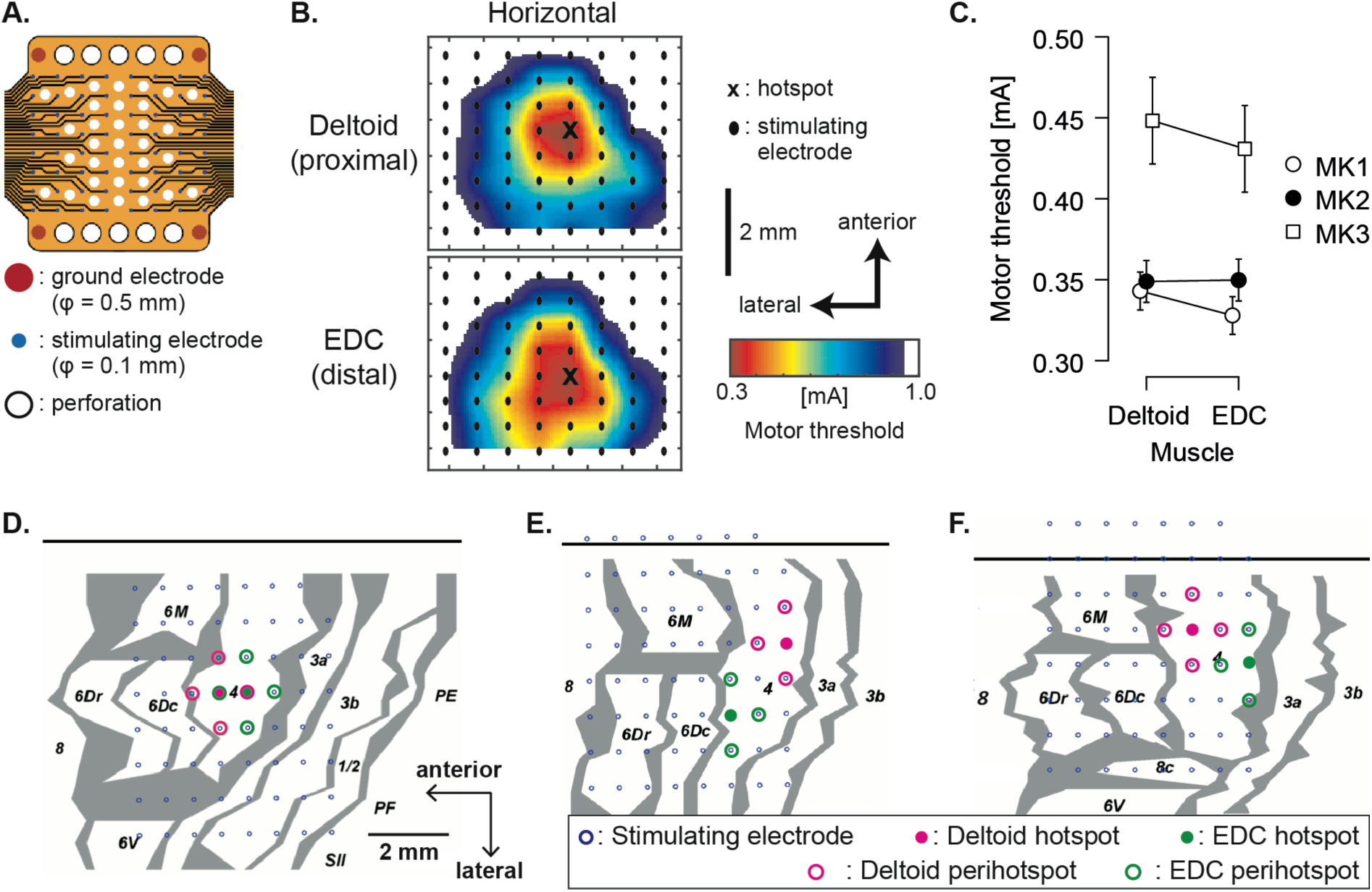
CSS electrode arrays and raw M1 mapping results. **(A)** The 64-channel microelectrocorticographic electrode arrays implanted on the left primary motor cortex (M1) for cortical stimulation mapping (CSS). **(B)** Motor maps of the right deltoid and extensor digitorum communis (EDC) muscles in the horizontal posture for marmoset MK1. In each map, the hotspot (black cross) denotes the electrode location with the lowest motor threshold (MT). Topographical maps were constructed using the average of 16 MT maps. Each map consisted of the MTs measured at all 64 electrode locations, which were spatially interpolated. **(C)** Hotspot MT of the deltoid and EDC muscles. Dots and error bars represent the mean and 95% confidence interval. **(D–F)** Locations of the stimulating electrodes and hotspots of deltoid and EDC muscles overlaid on the histological mapping results for the three marmosets MK1 **(D)**, MK2 **(E)**, and MK3 **(F)**. The numbers in the figures indicate the cortical areas, the top black horizontal line represents the interhemispheric fissure, and the gray areas represent the architectural borders. Hotspot locations were constant among the body postures.

**Figure 2.**
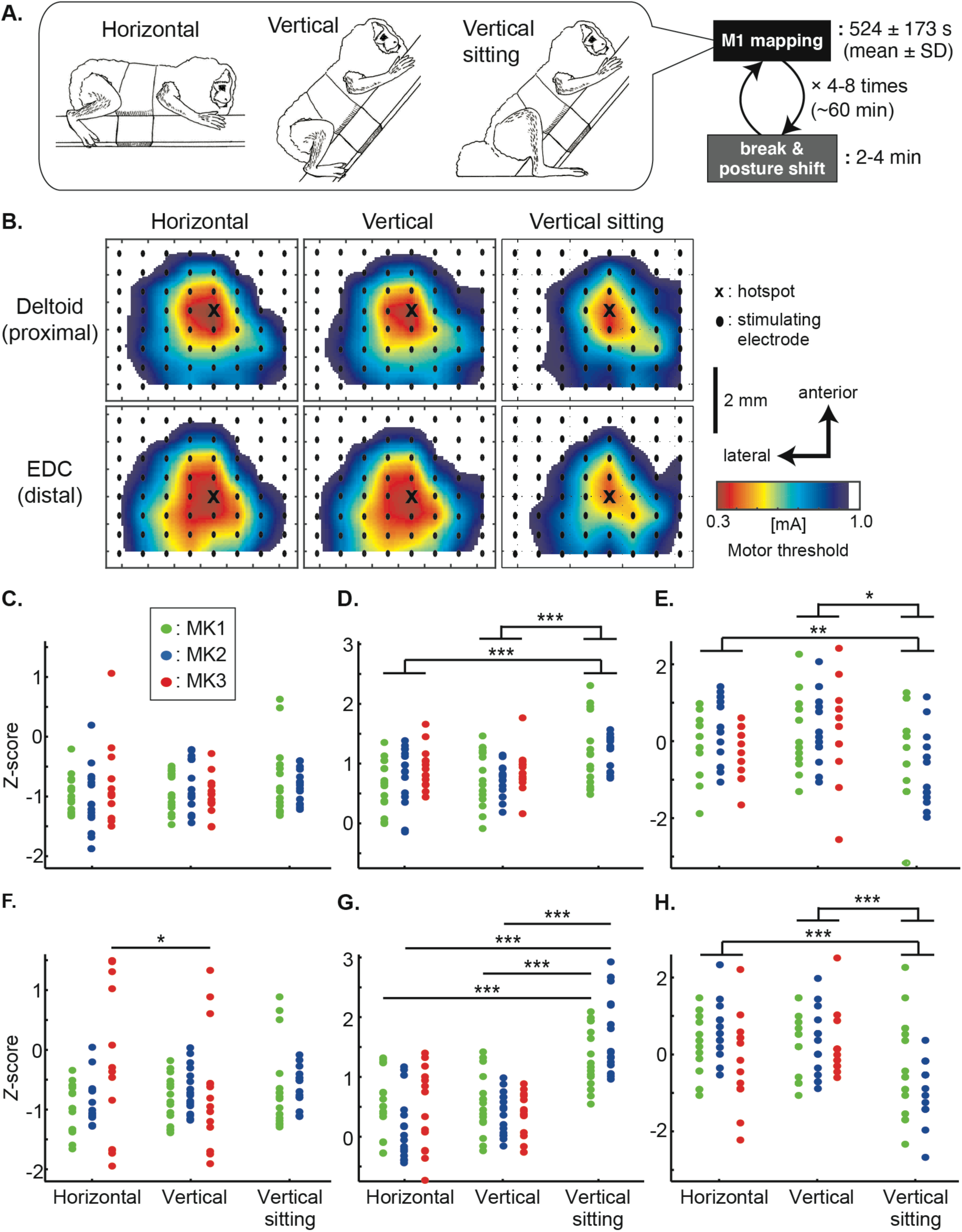
Experimental schema and modulation of forelimb muscle motor map in M1 by posture. **(A)** Procedure for a single M1 mapping session. We mapped forelimb motor representations in the M1 with the animal in one of three whole-body postures. Postures were tested in pseudorandom order. **(B)** Motor maps of the right deltoid and extensor digitorum communis (EDC) muscles in the three whole-body postures for marmoset MK1. **(C)** Hotspot MT of the deltoid muscle. **(D)** Perihotspot MT of the deltoid muscle obtained by averaging the MTs of electrodes over area 4 within a 1-mm surrounding the hotspot. The hotspot and perihotspot MTs were concatenated and z-scored separately in each monkey and muscle. **(F)** Hotspot MT and **(G)** perihotspot MT of the EDC muscle. **(E)** Map sizes of the deltoid and **(H)** EDC muscle representations. Map sizes reflect the number of electrodes in which MT was ≤0.65 mA, transformed into z-score separately for each monkey and muscle. *: p < 0.05, **: p < 0.01, ***: p < 0.001, by post-hoc pairwise comparisons using the Bonferroni procedure.

The use of nanomaterial coatings reduced electrode impedance and increased charge injection capacity, which ensured reliable stimulus current output for many weeks. Each stimulating electrode was 100 μm in diameter, and the distance between electrode pairs was 900 μm in the mediolateral direction and 700 μm in the anteroposterior direction. Four interconnected stimulation ground contacts (diameter, 500 μm) were located at the corners of the sheet.

### Surgical procedure

The surgical procedures for μECoG and electromyographic (EMG) electrode implantation were described previously (Tia et al. 2017; Kosugi et al. 2018). Briefly, anesthesia was induced by intraperitoneal injection of 0.05 mg/kg medetomidine, 0.5 mg/kg midazolam, and 0.5 mg/kg butorphanol. Atropine (0.10 mg/kg) and prednisolone (0.15 mg/kg) were injected intramuscularly (i.m.) immediately after anesthetic injection. During surgery, anesthesia was maintained with 1.5%–2.5% isoflurane inhalation, and the oxygen saturation level was continuously monitored. After the initial surgery, animals were administered the antibiotic cefmetazole (25 mg/kg, i.m.), the anti-inflammatory corticosteroid prednisolone (0.15 mg/kg, i.m.), and the analgesic mefenamic acid (6.5 mg/kg, oral administration) daily for 14 consecutive days. When intensive postsurgical care was required, the animals were anesthetized with isoflurane (0.5-3%) and injected subcutaneously with lidocaine for analgesia.

For implantation of the μECoG electrode array sheet, a 9 × 5 mm craniotomy was performed over the left hemisphere (coordinates relative to bregma: 0–9 mm anterior and 2–7 mm lateral). The μECoG electrode sheet was then laid onto the dura using a micromanipulator, and a piece of artificial dura mater was placed between the sheet and skull to hold the electrodes in place and prevent drying of the dura. A head chamber made of Ultem (height, 15 mm; width, 18 mm; length, 16 mm) was attached to the skull with stainless steel screws and dental acrylic to fix the electrode connectors. The area inside the chamber was washed with sterile saline every day for 7–10 days postoperatively to reduce reactive tissue formation around the electrode arrays and then filled with silicone polymer (Kwik-Cast, World Precision Instruments, Sarasota, FL).

The EMG electrodes were implanted 4–6 days before µECoG implantation surgery. Briefly, pairs of multistranded stainless steel wires (AS634, Cooner Wire, Chatsworth, CA) were implanted subcutaneously into four muscles of the right forelimb: deltoid, triceps brachii (TB), extensor carpi radialis (ECR), and extensor digitorum communis (EDC). Additional EMG wires were implanted into the biceps brachii and flexor digitorum superficialis muscles of MK3. All muscles were first located by anatomical features and identities verified by movements elicited in response to low-intensity electrical stimulation directly applied to the muscles. The electrode pairs were implanted approximately 5 mm apart.

At the end of the experiment, histological mapping was performed to identify the cortical regions covered by the µECoG array. Briefly, the monkeys were deeply anesthetized with ketamine (15 mg/kg) and pentobarbital (75 mg/kg), then perfused through the heart with phosphate-buffered saline and paraformaldehyde. A block of the postmortem cortex containing the stimulated areas was cut parasagittally, and 50-μm histological sections were mounted and Nissl stained. The location of the entire array over the cortex was then reconstructed from serial sections. Detailed criteria used to distinguish between premotor, primary motor, and somatosensory areas were described in our previous study (Tia et al. 2017). In brief, we followed recent references (Burish et al. 2008; Burman et al. 2008, 2014a, 2014b) and the marmoset three-dimensional digital brain atlas (Woodward et al. 2018) for histological analysis and identification of areas in the frontal cortex. Area 4 is characterized by the absence of layer IV and the presence of large pyramidal cells in layer V. Area 6Dc is distinguished by features similar to those of area 4, but with smaller cells in layer V. Area 6M is medial to 6Dc but in contrast to areas 4 and 6Dc includes an incipient layer IV and larger cells in layer V than area 6Dc. Area 3a consists of an identifiable layer IV and large cells in layer V, although these cells are smaller than those present in area 4. Histological borders were plotted as transition zones of various widths reflecting sources of uncertainty, such as test– retest variability (assessed by repeated plotting by the same observer on different days) and interference of histological artifacts.

### Cortical stimulation mapping

Cortical mapping of M1 was performed using a previously developed automatic cortical stimulation system (Takemi et al. 2017) and an algorithm for stochastic estimation of the motor threshold (MT; Kosugi et al. 2018), the stimulus intensity eliciting muscle twitch with 50% probability. The MT values determined at different points across the electrode matrix were then used to construct M1 maps (Fig. 1B). The stimulus train consisted of five biphasic pulses (250 µs cathodal and 250 µs anodal) delivered at 1,000 Hz. The stimulus current was generated by an isolated output source (SS-203J, Nihon Kohden, Tokyo, Japan) and controlled with an analog output module (NI PCIe-6321, National Instruments). The maximum stimulator output was set to 1.0 mA. The EMG signals evoked by cortical stimulation were band-pass filtered (1–2,000 Hz with 2nd order Butterworth filter) and digitized at 4,800 Hz using a bioamplifier (g.USBamp, g.tec medical engineering GmbH, Graz, Austria).

During M1 mapping, peak-to-peak EMG amplitudes of the deltoid, TB, ECR, and EDC muscles were continuously calculated over the last 80 ms. A peak-to-peak EMG amplitude exceeded 50 µV prior to stimulation onset (−80 ms to 0 ms) was considered voluntary muscle activity, and cortical stimulation was automatically stopped until termination of this activity. In addition, the peak-to-peak EMG amplitude elicited 10–20 ms following the first pulse of the stimulus train was calculated as the motor-evoked potential (MEP). Any MEP amplitude that exceeded a predetermined threshold was considered significant (i.e., a muscle twitch was evoked). We set this predetermined threshold to 60 µV, 1.2 times higher than the EMG amplitude considered indicative of voluntary muscle contraction, to minimize the detection of false-positive MEPs. The MT was separately but simultaneously estimated for all electrodes using the modified maximum likelihood algorithm (Kosugi et al. 2018), which fits the cumulative Gaussian distribution of measured MEP probability at different stimulation intensities. We first modeled the probability (*p*) of obtaining a significant MEP (<60 µV) at a particular stimulus intensity using a cumulative Gaussian. The log-likelihood of *p* after *n* stimulations was then calculated, and the stimulation parameters that maximized this log-likelihood were identified. The algorithm then adjusted the next stimulus intensity based on whether the present stimulus intensity elicited a significant MEP in the muscle being assessed. The MT estimation at a given electrode was terminated if either the last four stimuli at 1.0 mA did not evoke a significant MEP or if the changes in intensity at the electrode were within ±0.01 mA for the last four stimuli. The latter criterion was not applied until 20 stimuli were delivered by the electrode for reliable MT estimation (Kosugi et al. 2018).

The stimulation electrode was randomly chosen from among the 64 contacts in each trial and the initial stimulus intensity was set at 0.35 mA. The stimulus train was repeated every 300 ms but was not applied to the same electrode within a 2-s interval to avoid inducing cortical plasticity and to ensure good test–retest reliability of the mapping results (Nudo et al. 1990; Kosugi et al. 2018). This necessitated the insertion of pauses within the stimulation sequence as the mapping progressed and the MT was determined for most of the 64 electrodes. The M1 mapping system, including the MT estimation algorithm, was programed in MATLAB 2013a (MathWorks, Natick, MA).

### Experimental protocol

In the present study, we constructed M1 maps for only the deltoid and EDC (i.e., one proximal and one distal forelimb muscle) in the three whole-body postures, as our aim was to demonstrate the dynamicity of M1 muscle representations rather than the generalizability of this pattern over the entire forelimb. Moreover, restricting mapping to two muscles reduced the experiment duration and facilitated animal cooperation.

The M1 maps in all three postures were constructed for MK1 and MK2 over 8 and 4 weeks, respectively. In total, 16 M1 maps were acquired for each posture and muscle per monkey. For MK3, M1 maps were obtained in horizontal and vertical postures over a period of 6 weeks, and 14 M1 maps were acquired for each posture and muscle. Of note, the vertical-sitting posture was considered based on examination of horizontal and vertical posture results for MK3 and was included only in MK1 and MK2. Unfortunately, testing in the vertical-sitting was not feasible for MK3 due to imperative randomization of the order of postural conditions to mitigate potential biases associated with changes in electrode impedance.

For the M1 mapping session, the marmosets were jacketed, and the jacket was fixed to a wooden pole (diameter, 3.5 cm) covered by a urethane sheet (thickness, 5 mm) with marks where the animals’ limbs were to be placed. The head position was fixed by clamping the head chamber with screws. In all postures, head orientation was the same as that of the trunk (forward and upward as in Fig. 2A). However, visual inputs differed between the horizontal and vertical/vertical-sitting conditions. In the horizontal posture, the animals’ field of view included the fixation pole and the wall of the experiment room, whereas in the vertical and vertical-sitting postures, the visual field comprised the pole, wall, and ceiling of the experiment room. We ensured that forelimb posture and skin afferent were controlled and constant across conditions (i.e., flexed and in contact with the pole). Monkeys were habituated to this experimental condition by progressively increasing restraint duration to 75 min over a month, with food rewards provided every 8–16 min. During the actual experiment, the monkey was positioned as shown (Fig. 2A), the µECoG electrodes were connected to the cortical stimulator, and M1 stimulation mapping was initiated. During stimulation, we visually monitored the position of the limbs. Cortical stimulation was automatically stopped if the marmoset moved the right forelimb and manually stopped if the other limbs moved. In both cases, we manually repositioned the limb to its initial mark.

In a single session (lasting approximately 60 min), we performed four to eight M1 mappings depending on the condition of the animal (Fig. 2A). Descriptive statistics of the duration (mean ± SD) of individual mappings are summarized in Table 1. We did not start a new mapping if the session exceeded 1 h. The session was also terminated if the animals moved their limbs frequently. The marmosets were given breaks between mappings during which they were rewarded with sweets, and posture was adjusted if needed. The order of test muscles and postures was pseudorandomized across sessions to minimize the effects of restraint time.

**Table 1.** Duration of cortical stimulation mapping in different body postures.

|  | Horizontal [s] | Vertical [s] | Vertical-sitting [s] |
| --- | --- | --- | --- |
| MK1 | 430 ± 128 | 493 ± 186 | 446 ± 111 |
| MK2 | 495 ± 186 | 614 ± 168 | 599 ± 227 |
| MK3 | 573 ± 174 | 554 ± 180 | Not tested |

Values represent the mean ± SD.

### Data analysis

To evaluate the flexibility of topographical motor representations in M1, we calculated four functional indices for each map: map size, hotspot location (for the test muscle), hotspot MT, and perihotspot MT. The map size was calculated as the number of electrodes over area 4 for which the MT was lower than 0.65 mA to ensure test–retest reliability (Kosugi et al. 2018). The hotspot location was defined as the position with lowest MT among the 64 electrodes, and the perihotspot MT was defined as the mean for all electrodes over area 4 within 1 mm surrounding the hotspot.

Each map was first z-transformed to allow comparisons of the aforementioned indices between monkeys, muscles, and postures. An analysis of variance (ANOVA) was independently applied to the z-scores of deltoid and EDC map sizes with monkey (MK1, MK2, MK3) and posture (horizontal, vertical, vertical-sitting) as between-subjects factors. The hotspot MT and perihotspot MT were calculated separately for each monkey and muscle, then concatenated and transformed into z-scores. To test whether body posture modulated different aspects of the functional M1 representation, an ANOVA was independently applied to the z-scores of deltoid and EDC MTs, with monkey and posture as between-subjects factors and location (hotspot and perihotspot) as a within-subject factor. Post-hoc t-tests were performed using the Bonferroni procedure to correct for multiple comparisons. The type I error was set to 0.05 and all tests were conducted using IBM SPSS Statistics, version 22 for Windows.

We also analyzed two additional parameters that could influence mapping results: prestimulus muscle activity and mapping duration. In the current study, cortical stimulation automatically stopped when EMG activities within 80 ms before stimulus onset exceeded 50 µV (as this was considered to reflect voluntary movement as noted). However, a previous study in humans suggested that voluntary muscle contraction could affect corticospinal output profiles for up to 160 ms after movement (Chen et al. 1998). Thus, we compared the average rectified EMG amplitudes in the 80–160 ms window preceding cortical stimulation among different postures. An ANOVA was separately applied to prestimulus activities of the deltoid, TB, ECR, and EDC muscles, with monkey and posture as between-subjects factors. In addition, prolonged mapping duration can induce central fatigue, which reduces the descending corticospinal volleys evoked by cortical stimulation to M1 (Brasil-Neto et al. 1993) and thus increases MT. To test whether mapping duration affected MT, we analyzed the correlation between mapping duration and hotspot MT z-scores of the deltoid and EDC muscles for each monkey.

## Results

### Deltoid and EDC hotspot MTs and locations

A two-way ANOVA was performed to compare the MT values between deltoid and EDC muscles, using muscle as the within-subject factor and monkey as the between-subject factor, which revealed no significant main effect of muscle (F_(1, 121)_ = 2.62, p = 0.108) and no significant muscle × monkey interaction (F_(2, 121)_ = 0.86, p = 0.426), indicating no discernible difference in the MT values between deltoid and EDC muscles (Fig. 1C). In addition, the hotspots of the deltoid and EDC muscles were found at different electrode locations in all the three marmosets (0.7 mm apart in MK1, 2.3 mm in MK2, and 1.7 mm in MK3) (Figs. 1D-F). Furthermore, these hotspots were consistently observed over area 4 regardless of body posture (Fig. 2B).

### Modulation of perihotspot MTs and map size by posture

The motor representations of the forelimb muscles in the M1 were modulated mostly by their hindlimb posture and not by the orientation of their body axis relative to the gravitational vertical line. This modulation was observed for the proximal and distal forelimb muscles, occurring primarily in the perihotspot area.

An ANOVA with z-scored deltoid muscle MT (Figs. 2C,D) as the dependent variable demonstrated an interaction between posture and location (F_(2, 116)_ = 4.44, p = 0.014); however, there was no interaction between posture, location, and monkey (F_(3, 116)_ = 2.40, p = 0.071). Post-hoc pairwise comparisons revealed that MT in the perihotspot area was higher in the vertical-sitting posture than in the horizontal and vertical postures (p < 0.001). No difference was found in perihotspot MTs between the horizontal and vertical postures. Furthermore, the hotspot MTs were not altered among the different postures.

An ANOVA with z-scored map size of the deltoid muscle (Fig. 2E) as the dependent variable revealed a main effect of posture (F_(2,116)_ = 8.39, p < 0.001); however, no interaction was observed between posture and monkey (F_(3, 116)_ = 1.53, p = 0.21). Post-hoc analysis revealed that the map size was smaller in the vertical-sitting posture than that in the vertical (p = 0.001) and horizontal postures (p = 0.035). No difference was present between the horizontal and vertical postures.

Posture-dependent changes in the MTs of the EDC muscle were almost identical to those of the deltoid muscle; however, they were slightly inconsistent between monkeys. An ANOVA with z-scored EDC muscle MT (Figs. 2F,G) as the dependent variable revealed an interaction between posture, location, and monkey (F_(3, 116)_ = 5.24, p = 0.002). Post-hoc pairwise comparisons revealed that in MK1 and MK2, the perihotspot MT was higher in the vertical-sitting posture than in the horizontal and vertical postures (p < 0.001). No difference was found in the perihotspot MTs between the horizontal and vertical postures. Furthermore, hotspot MTs were not altered among the different postures. Conversely, in MK3, pairwise comparisons revealed that the vertical and horizontal postures significantly modulated the hotspot MT (p = 0.042) but not the perihotspot MT (p = 0.70).

ANOVA using the z-scored map size of the EDC muscle (Fig. 2H) as the dependent variable indicated a main effect of posture (F_(2, 116)_ = 11.5, p < 0.001); however, there was no indication between posture and monkey (F_(3, 116)_ = 2.32, p = 0.079). Post-hoc analysis revealed that map size was smaller for the vertical-sitting posture than that for the vertical and horizontal postures (p < 0.001). No difference was detected between the horizontal and vertical postures.

### Changes in MT over the experimental period

To examine the day-to-day variations in MTs, we constructed graphs with dates on the horizontal axis and z-scored MTs on the vertical axis (Fig. 3), which displays the same daily sessions on the same x-coordinate and enables a clear observation of within-day MT variability and longitudinal changes. Our regression analysis revealed statistically significant positive relationships between MTs and the number of days after ECoG implantation in most instances, indicating a constant rise in the minimum current intensity required to ignite corticomotor neurons beneath the array electrodes, possibly because of the growth of dura or scar tissue.

**Figure 3.**
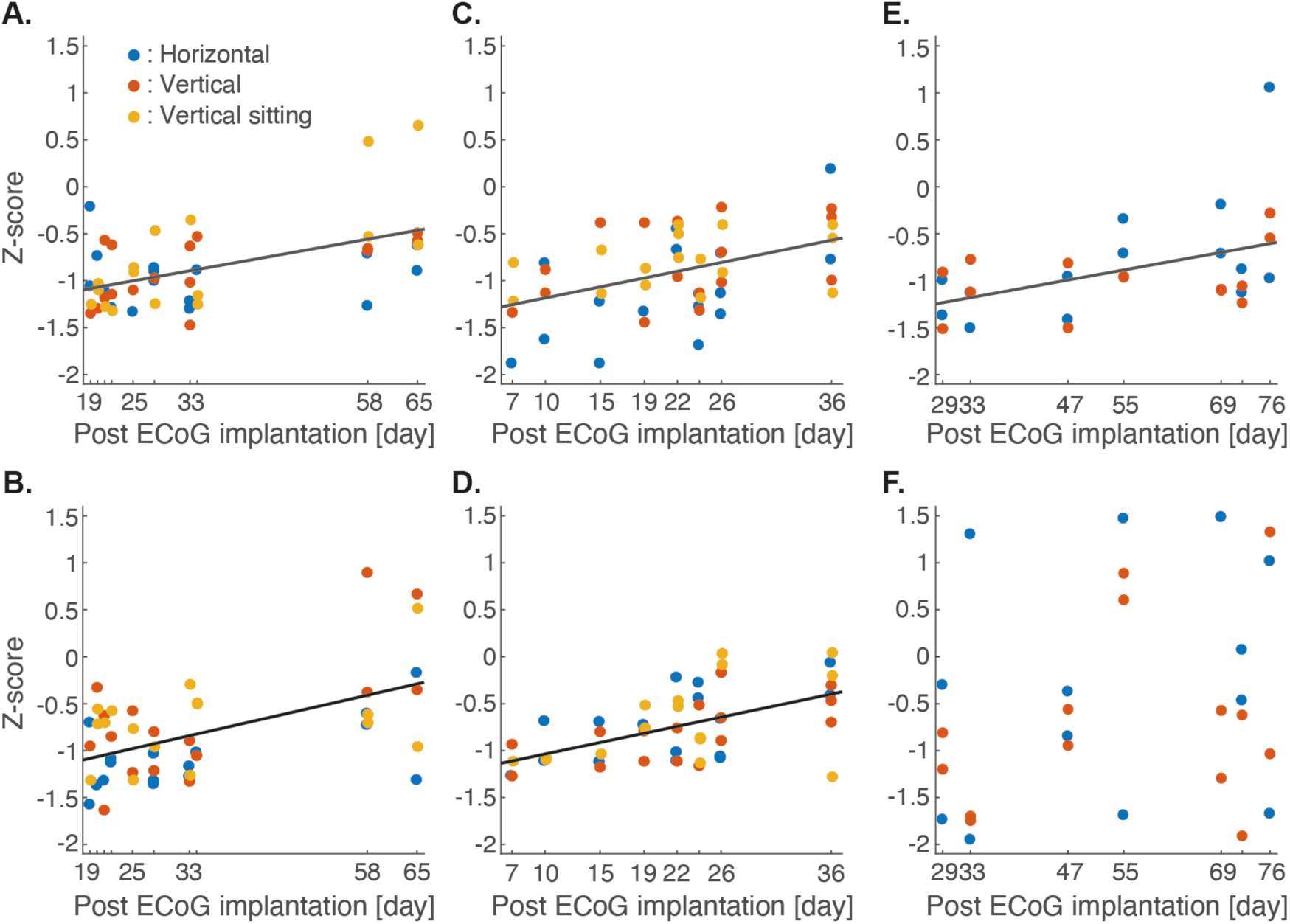
Progressive increases in motor threshold with time after electrode implantation. **(A–F)** Day-to-day variation in hotspot MT values for the deltoid muscle **(A, C, E)** and EDC muscle **(B, D, F)** for marmosets MK1 **(A, B)**, MK2 **(C, D)**, and MK3 **(E, F)**. The MT values were z-scored for each marmoset and muscle. Linear trend lines were independently estimated for each plot and incorporated into the respective panels if the null hypothesis that the slope coefficient is zero was statistically rejected (p < 0.05).

### Mapping duration

Table 1 summarizes the duration of cortical stimulation mapping. An ANOVA with mapping duration as the dependent variable demonstrated no main effect of posture (F_(2, 116)_ = 1.30, p = 0.276) and no interaction between posture and monkey (F_(3, 116)_ = 1.10, p = 0.353) but a significant main effect of monkey (F_(2, 116)_ = 6.70, p = 0.002). There were no correlations between the z-scored mapping duration calculated separately for each monkey and muscle hotspot MT (deltoid: r_(122)_ = 0.093, p = 0.31; EDC: r_(122)_ = 0.089, p = 0.32).

### Prestimulus muscle activity

In all four muscles, the average rectified EMG amplitudes in the 80–160 ms window preceding cortical stimulation were generally small (<10 µV) and thus unlikely to reflect voluntary contraction (Table 2). According to ANOVA, there were no main effects of posture on prestimulus deltoid, TB, and ECR muscle activities (F_(2,116)_ = 0.54–1.06, p = 0.35–0.58) and no interaction between posture and monkey (F_(3, 116)_ = 0.38–1.81, p = 0.15–0.77). However, ANOVA demonstrated a posture × monkey interaction on prestimulus EDC muscle activity (F_(3, 116)_ = 4.11, p = 0.008), and post-hoc analysis revealed that prestimulus EDC muscle activity in MK3 was larger during the horizontal posture than the vertical posture (p = 0.001). This difference may account for the posture-dependent modulation of EDC hotspot MTs observed in MK3 (Fig. 2G). In contrast, prestimulus EDC muscle activity did not differ among the postures in MK1 and MK2.

**Table 2.**
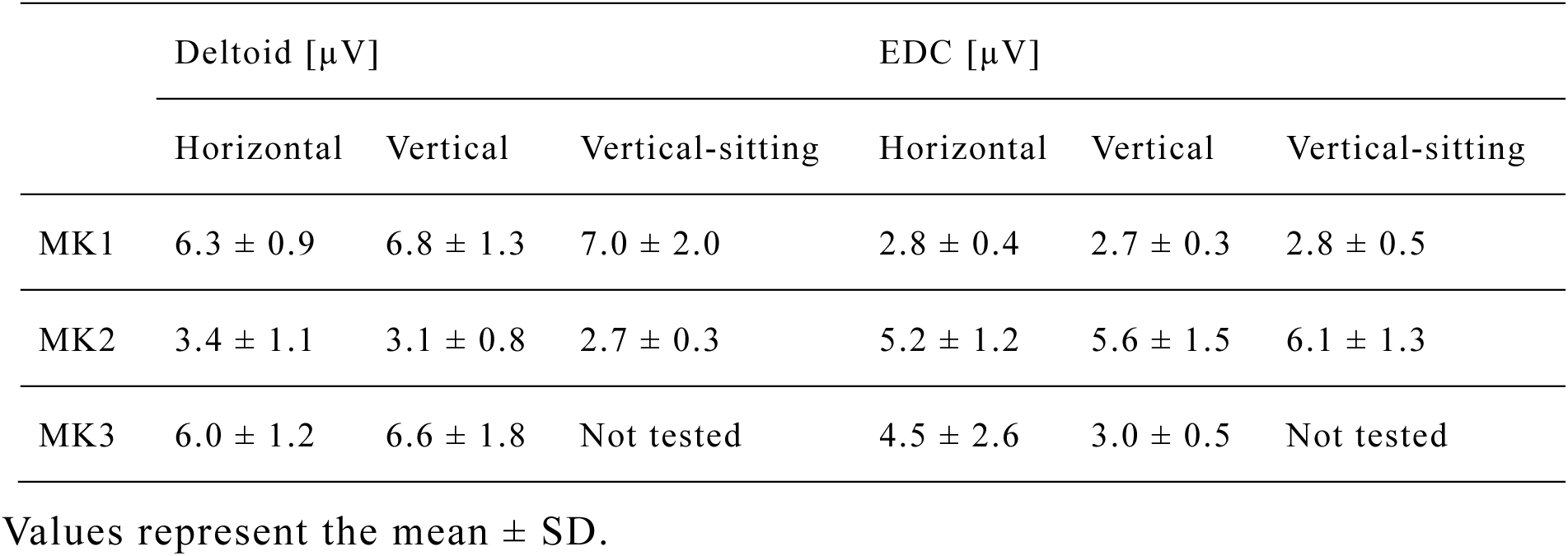
Averaged rectified EMG amplitudes of the deltoid and EDC muscles for the 80–160-ms window preceding cortical stimulation.

## Discussion

### Sensorimotor inputs and dynamic M1 motor maps

The findings of this study serves as a proof of concept, revealing that a rapid stimulation mapping system with chronically implanted cortical electrodes can capture the dynamic regulation of M1 forelimb motor maps under natural conditions. The findings showed differences in the perihotspot MT, a biomarker of M1 motor maps, between vertical and vertical-sitting postures of the hindlimbs although the animals’ visual input and body orientation were identical. No such difference was observed between the vertical and horizontal postures despite different visual field and vestibular inputs. The modulation of M1 maps in the perihotspot region but not in the hotspot regions suggests the involvement of specific neuronal populations, excluding the largest and most excitable neuronal pools projecting to the forelimb muscles. Moreover, the consistency in mapping duration across postures excluded biases associated with central fatigue on motor representations (Brasil-Neto et al. 1993). While our primary focus was to demonstrate the detectability of rapid changes in M1 maps, our findings provide insights into potential neurophysiological mechanisms, as discussed below.

The observed increase in MT and decrease in map size in the vertical-sitting posture compared with that in the vertical posture indicate a potential role of somatosensory inputs from the hindlimbs in modulating M1 forelimb representation. These alterations may occur because of the (1) an inhibition of corticomotor neurons for the deltoid and EDC muscles and (2) an inhibition of the motoneuron pools at the spinal level. The first scenario could involve afferent signals from the hindlimb region of somatosensory areas 3a/3b. In marmosets, cortical projections to the M1 area innervating forelimb muscles primarily arise from the dorsocaudal and medial premotor cortex, the primary and secondary somatosensory areas, and a subdivision of the posterior parietal area—termed PE (Burman et al. 2014a). Among these areas, fibers from areas 3a and 3b have more connections to the pericenter of forelimb M1 representation than to the center (i.e., the general location of the hotspot; Burman et al. 2014a). Area 3b receives information from cutaneous mechanoreceptors, some of which innervate the glabrous skin on the foot sole. Area 3a receives input from the muscle spindles, which code muscle stretch information, as well as joint receptors (Huffman and Krubitzer 2001). Thus, if cortical inputs from areas 3a and 3b influence forelimb motor representations, this modulation should coincide with changes in cutaneous feedback and muscle length, as supported by our current results.

Another interpretation of our results is that the dynamic regulation of motor representations is attributable to the spinal level, evincing that postural variations, such as changes in the joint angle (Chapman et al. 1991; Knikou and Rymer 2002), and input from cutaneous mechanoreceptors considerably influence intralimb spinal excitability (Knikou and Conway 2001; Zehr et al. 2014). This phenomenon may extend to distant body segments, whereby the peripheral stimulation of one limb induces reflex motor activity in the contralateral limb or in a remote body part (Nakajima et al. 2013; Butler et al. 2016). Interlimb reflexes involve propriospinal neurons that project through multiple segments of the spinal cord, contributing to context-dependent forelimb and hindlimb coordination (Nakajima et al. 2013; Hurteau et al. 2018; Shepard et al. 2021). In our experiment, modifying the peripheral inputs to the hindlimbs in the sitting posture could inhibit the excitability of the forelimb motoneuron pools. This aspect warrants further investigation, particularly considering the reported interspecies differences in the propriospinal transmission of corticospinal excitation to motoneurons, which is associated with locomotor mode (e.g., quadrupedal/bipedal and arboreal/terrestrial), hand dexterity, and other biomechanical specializations (Lemon 2008).

In addition to sensory inputs, the modulation of deltoid and EDC representations in M1 might stem from the contraction of muscles in remote body segments, such as trunk and hindlimb, where EMG signals were not measured in our experiment (Tazoe et al. 2009; Kato et al. 2016; Sasaki et al. 2018). The cortical or spinal origin of these remote effects is debatable. At the cortical level, prior studies reported that muscle contraction can influence the motor representation of distant body segments by modulating intracortical inhibitory circuits (Tazoe et al. 2007; Kato et al. 2016) and facilitating the spread of neuronal activity to nearby cortical loci (Capaday et al. 2011). Conversely, at the spinal level, evidence suggests that the contraction of contralateral muscles can alter the excitability of ipsilateral motoneuron pools through interneuronal circuits (Stedman et al. 1998; Hortobágyi et al. 2003; Bunday and Perez 2012).

### Limitations and perspectives

Our study revealed the dynamic features of M1; however, there were certain inherent limitations that warrant careful consideration. Examining only one proximal and one distal forelimb muscle may limit the generalization of our findings to the entire forelimb, as the influence of hindlimb posture on the representation of the other forelimb muscles remains unexplored. Although we considered the effect of hindlimb sensory input on forelimb muscle corticospinal excitability, the specific contributions of skin afferents, muscle proprioceptors, and joint proprioceptors alongside the impact of hindlimb muscle tone remain indistinguishable. The neural pathways underlying our observations—whether cortical, downstream, or a combination of both—remain speculative.

To enhance our understanding, future studies could rigorously manipulate hindlimb sensory input through various means, such as skin stimulation, footpad pressure alterations, or pharmacological blockade of skin afferents, thus enabling a clearer delineation of the roles played by tactile and proprioceptive inputs (e.g., muscle length and joint angle). Additionally, assessing EMG activity in remote body parts, particularly postural muscles, can provide valuable insights into potential differences between various postures. The simultaneous measurements of cortical and spinal excitability coupled with pharmacological blockade techniques can further elucidate the contributions of various neuronal pathways (Jacobs and Donoghue 1991; Adkins et al. 2006).

A technical limitation of our study lies in the characteristics of CSS. While we have previously demonstrated that CSS and ICMS can yield comparable and statistically correlated motor maps (Takemi et al. 2017), the higher current intensity requirement of CSS (Burish et al. 2008; Cooke et al. 2012) and its potential for trans-synaptic activation of layer 5 pyramidal neurons via the axonal excitation of upper layer cortical interneurons (Nowak and Bullier 1998; Wongsarnpigoon and Grill 2012) introduce challenges. CSS lacks the precision of ICMS and can indirectly activate distant cortical and subcortical structures through horizontal connections, rendering it difficult to directly infer physiological insights from ICMS literature and necessitating caution in interpretation. Measuring and estimating the extent of current spread and trans-synaptic excitation is crucial for advancing the application of CSS.

The electrode array density (700–900 µm, anteroposterior × mediolateral) imposes another limitation. We specified the present interelectrode distance by referring to a study demonstrating the optimal spacing of cortical surface electrode arrays (Slutzky et al. 2010). However, given that M1 forelimb representation in marmosets only extends up to ∼2 × 3 mm (anteroposterior × mediolateral; Burish et al. 2008), no more than 16 electrodes could have been in this area. This relatively low spatial resolution might affect the observations in this study, although our mapping procedure could still discriminate the hotspot locations of the deltoid and EDC muscles and capture the dynamic spatial patterns of motor representations.

### Implications: neuromodulation through body posture manipulation

Inferring the functional significance of our findings with regard to marmoset motor behavior poses some challenges. A previous study has reported that body posture affects hand use in marmosets (Hashimoto et al. 2013). Accordingly, our study suggests that afferent signals conveying information regarding the whole-body posture can modulate motor maps by altering the area and excitability profile of cortical motor representations. We posit that whole-body posture can serve as a modulator of M1 excitability, similar to human studies reporting differences in resting-state brain activity between upright, supine, and sitting postures (Thibault et al. 2014; Spironelli et al. 2016). This assumption carries practical implications for motor skill learning, rehabilitation following brain injuries, and basic M1 mapping experiments.

We presumed that practicing motor skills in different body postures, particularly those enhancing neuronal activity in motor representations associated with skilled movements, can facilitate the learning process. Reportedly, the artificial upregulation of M1 excitability, such as via intermittent theta burst stimulation, can accelerate motor skill learning (Teo et al. 2011). The application of intermittent theta burst stimulation to the ipsilesional M1 hand area following a hemiparetic stroke promotes the recovery of upper limb motor function (Chen et al. 2019). Thus, conducting motor rehabilitation in postures that increase neuronal excitability in the cortical motor representations governing paralyzed extremities can enhance the recovery process.

It is also important to emphasize the necessity of conducting all motor mapping experiments in the same body posture within a study, especially when investigating the reorganization of the topographical patterns of motor representations. This ensures the dissociation of changes in the M1 maps driven by long-term reorganization due to interventions such as motor training, limb amputation, and cortical damages (Nakagawa et al. 2020) from those resulting because of short-term alterations in afferent inputs due to postural changes. Furthermore, as neurons rapidly adjust their sensitivity to afferent inputs in response to alterations in cognitive demands and behavioral states (e.g., sleeping, awake resting, or during tasks; Ferguson and Cardin 2020), the motor map, reflecting the level of neuronal excitability in M1, possibly represents not only simple motor parameters (e.g., skeletal muscles or movement directions) but also cognitive, mental, and circadian states (Levinthal and Strick 2012; Dum et al. 2019). Future studies on motor control and neuroplasticity need to consider the complex overlapping organization of neural representations in M1.

## Acknowledgement

We thank Dr. Yumiko Yamazaki, Mr. Masakado Saiki, and Mr. Masayuki Inada for their technical assistance. We thank Dr. Miki Taoka for his helpful advice on the histological analysis. This work was supported by the Brain/MINDS project from AMED Japan to AI. MT had been supported by JST PRESTO (#JPMJPR18J6).

